# InVitroGap: an open-source tool for automated quantification of wound closure in the in vitro scratch assay

**DOI:** 10.64898/2026.06.19.733445

**Authors:** Rakesh K. Arya, Mohit Sindhani, Sahab Ram Dewala, Christopher J. Weight, Laura Bukavina

## Abstract

**Background:** Scratch assays are widely used to study wound closure in vitro, but quantitative image analysis remains constrained by manual variability, proprietary workflows, and tools requiring programming expertise. We developed InVitroGap, a Python-based application with a browser-accessible interface for automated quantification of scratch assay closure from sequential microscopy images.

**Methods:** RCC-ER and Renca cells were seeded in 96-well ImageLock plates and scratched using a WoundMaker device for uniform linear wounds or a 200 µL pipette tip for crisscross wounds. Phase-contrast time-lapse images acquired at 0, 24, and 48 h with an IncuCyte SX5 system were independently analyzed using IncuCyte 2023A Rev2 and InVitroGap. The InVitroGap pipeline combines Gaussian smoothing, gradient-based texture mapping, adaptive percentile thresholding, and morphological post-processing to quantify wound confluence and relative wound density (RWD). Agreement was evaluated using paired comparisons, Pearson and Spearman correlations, Bland-Altman analysis, and mean absolute error (MAE).

**Results:** InVitroGap measurements closely tracked IncuCyte outputs across both cell lines, with no significant between-method differences (p > 0.05), strong pooled correlations (R² = 0.964 for RWD; R² = 0.983 for wound confluence), and small mean biases (absolute bias ≤ 1.64%). The tool successfully processed crisscross wounds from brightfield image series, and a complete four-timepoint series was analyzed in approximately 10 seconds, with robust performance across distinct cell morphologies and wound geometries.

**Conclusions:** InVitroGap provides a transparent, computationally efficient, and platform-independent alternative for scratch assay analysis, delivering performance comparable to commercial systems while remaining freely accessible at https://invitrogap.vercel.app/.

**Highlights:**

- Open-source Python tool for automated, platform-independent in vitro scratch assay analysis
- Texture-based adaptive pipelines enable robust wound segmentation across cell types
- Quantifies wound confluence and relative wound density from time-lapse images
- Strong agreement with IncuCyte measurements in the tested datasets

## 1. Introduction

Cell migration is a fundamental biological process underpinning development, tissue repair, immune responses, and pathological conditions such as cancer metastasis and chronic inflammation[1, 2]. In vitro models, including the scratch assay (also known as the wound-healing assay), remain the most widely adopted means to study collective cell movement in a controlled experimental setting[3]. The assay is performed by establishing a confluent monolayer of adherent cells, introducing a linear cell-free gap (“scratch”), and monitoring the migration of cells into this gap over time[4]. By quantifying the rate of gap closure, researchers can assess migratory potential, evaluate the effects of pharmacological agents and growth factors, and investigate molecular mechanisms governing cell–cell and cell–matrix interactions[5].

Despite its widespread adoption, quantitative analysis of scratch-assay images remains a significant methodological bottleneck. Manual approaches, most commonly performed using ImageJ, require users to delineate wound boundaries by hand, yielding workflows that are time-intensive, operator-dependent, and prone to inter-user variability [6–8]. To address these limitations, a range of automated tools spanning open-source software, cloud-based platforms, and deep-learning models have been developed, each with distinct capabilities and constraints. Among open-source tools, TScratch employs curvelet-based image segmentation to quantify open wound areas [9], while PyScratch provides a Python-based graphical user interface (GUI) for non-programmers[10]. The recently described CSMA tool offers ImageJ-compatible wound analysis [11], and deep-learning approaches utilizing U-Net architectures have also been explored for automated wound segmentation[12]. Although these tools have improved analytical objectivity, they can remain sensitive to imaging artifacts, may require non-trivial parameter tuning, or produce metric outputs that are not directly comparable to commercial platform standards, complicating cross-study interpretation. Cloud-based services such as Wimasis (WimScratch) enable scratch assay analysis via web image upload, but remote processing introduces data-transfer requirements that raise concerns about the handling of unpublished datasets, and limited user control over segmentation parameters can reduce reliability [13]. More recently, CLYTE Soφ 3.0 (Scratch Analyzer) has emerged as a web-based endpoint analysis tool; however, its workflow is restricted to pairwise comparison of two timepoints (initial versus final), precluding full time-course kinetic analysis, and its fixed processing pipeline offers limited parameter adjustability.

Commercial systems, notably the IncuCyte live-cell imaging platform (Sartorius), provide real-time automated analysis that generates validated metrics including wound confluence and relative wound density (RWD) [14, 15]. Similarly, the HoloMonitor platform (Phase Holographic Imaging) enables label-free wound healing quantification, but requires the proprietary HoloMonitor instrument and App Suite, restricting its use to laboratories equipped with that hardware. These platforms represent the current analytical gold standards; however, they are cost-prohibitive and strictly hardware-dependent, their proprietary “black-box” algorithms prevent methodological transparency and limit re-analysis of legacy datasets, and they cannot be applied to images acquired on alternative microscope systems[16]. Collectively, this landscape spanning hardware-locked commercial systems, cloud-based services with data-transfer requirements, endpoint-only web tools, and open-source tools with limited metric standardization, reveals a persistent gap: the absence of a freely accessible, platform-independent tool that supports full time-series kinetic analysis, produces commercially validated metrics, preserves data locality, and offers user-adjustable segmentation without requiring programming expertise.

A workflow-level comparison of InVitroGap with representative commercial, cloud-based, and open-source scratch-assay image-analysis solutions is provided in Supplementary Table 1.

To address this gap, we developed InVitroGap, an open-source, standalone Python application accessible at https://invitrogap.vercel.app/. InVitroGap implements experiment-adaptive image-processing pipeline integrating Gaussian smoothing, Sobel gradient-based texture extraction, and locally referenced adaptive percentile thresholding to enable robust wound segmentation across diverse imaging conditions. Through an intuitive GUI, the application automates time-series analysis of sequential images from a single wound area, outputs standard quantitative metrics, wound confluence, RWD, and closure rate and completes a four-timepoint series in approximately 10 seconds, eliminating the need for programming expertise or proprietary hardware.

In this study, we describe InVitroGap’s design and algorithmic workflow, validate its accuracy against IncuCyte 2023A Rev2 software using identical phase-contrast datasets from human (RCC-ER) and murine (Renca) renal carcinoma cell lines, and demonstrate its robustness across non-standard wound geometries and imaging modalities. Our results establish InVitroGap as a reliable, transparent, and freely accessible tool for standardized quantification of collective cell migration in cancer biology and broader biomedical research.

## 2. Materials and Methods

### 2.1. Cell culture and wound-healing assay

Human renal clear cell carcinoma cells (RCC-ER) and murine renal adenocarcinoma cells (Renca) were kind gifts from Dr. Philip Abbosh MD, PhD, Fox Chase Cancer Center, PA, USA. Both cell lines were authenticated by short tandem repeat (STR) profiling and confirmed negative for mycoplasma contamination prior to use. RCC-ER cells were cultured in RPMI-1640 medium (Gibco, Thermo Fisher Scientific, Waltham, MA, USA) supplemented with 10% fetal bovine serum (FBS), 1% sodium pyruvate, and 1% penicillin–streptomycin. Renca cells were maintained under identical conditions with the addition of 0.1% non-essential amino acids. All cultures were maintained at 37 °C in a humidified atmosphere with 5% CO₂.

Cells were seeded in 96-well ImageLock plates (Sartorius, Göttingen, Germany) at densities of 30,000 cells per well (RCC-ER) and 37,000 cells per well (Renca) and allowed to reach near-confluence within 24 h. Uniform linear scratch wounds (∼700–800 µm wide) were generated using the WoundMaker tool (Sartorius). To evaluate software performance on non-standard wound geometries, crisscross scratches were independently generated using 200 µL pipette tips. Following scratch creation, all wells were washed twice with complete medium to remove cellular debris and replenished with 200 µL of fresh complete medium to assess baseline cell migration.

### 2.2. Image acquisition and data source

Following scratch creation, plates were transferred to the IncuCyte® SX5 live-cell analysis system (Sartorius) for automated imaging under physiological conditions (37 °C, 5% CO₂). Phase-contrast images were acquired using a 10× objective at fixed field positions every 24 h over a total duration of 72 h (time points: 0, 24, 48, and 72 h). Raw image files were exported in TIFF format for offline analysis. For the validation study, identical image sets were analyzed independently and in parallel by both the IncuCyte 2023A Rev2 commercial software and the InVitroGap application, enabling direct well-by-well comparison of wound-closure metrics. For figure presentation, representative cell-line-specific image panels and summary plots in Figures 3 and 4 were limited to 0, 24, and 48 h to preserve visual clarity, whereas the full 0–72 h dataset was retained for the complete time-course illustration in Figure 2 and for pooled agreement analyses in Figures 5 and 6. Brightfield images of crisscross wounds acquired at 10× using a standard laboratory microscope (EVOS XL Core, Thermo Fisher Scientific) were included to assess InVitroGap performance under non-IncuCyte imaging conditions; this auxiliary feasibility dataset was acquired through 48 h and is therefore presented to that endpoint in Supplementary Figure S1.

**Figure 1.**
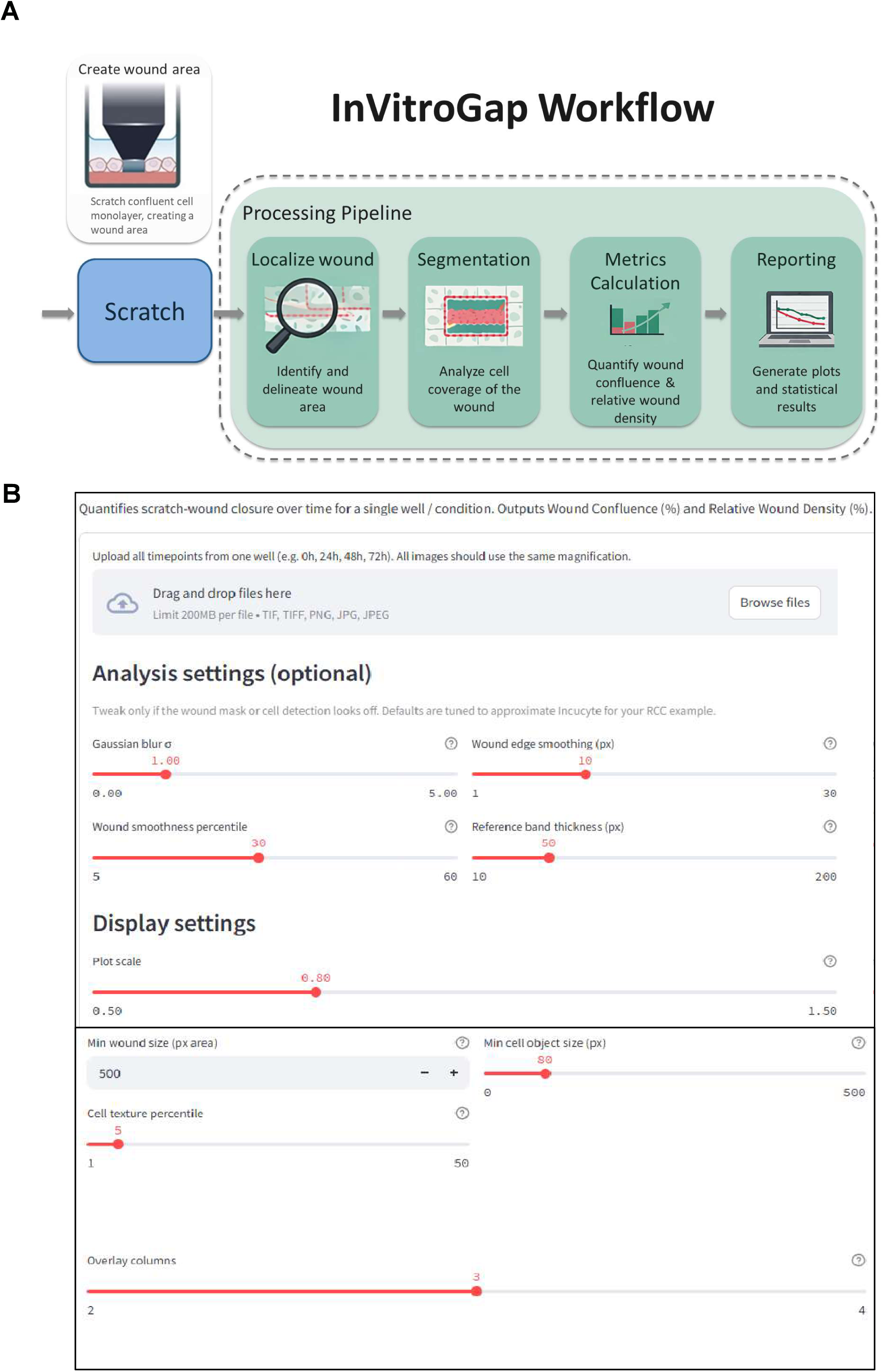
InVitroGap architecture and graphical user interface. **(A)** Schematic of the experiment-adaptive image-processing pipeline, illustrating sequential steps: Gaussian smoothing (σ = 1.0), Sobel gradient-based texture extraction, fixed wound ROI definition at t = 0 h, reference-band generation by dilation (50 pixels), and dynamic percentile thresholding for adaptive segmentation at subsequent time points. **(B)** Screenshot of the browser-accessible GUI showing drag-and-drop image upload and adjustable analysis parameters (Gaussian blur σ, wound smoothness percentile, reference-band thickness, minimum wound and cell-object sizes, cell-texture percentile). Default parameter values are pre-configured to facilitate use by investigators without image-processing expertise.

**Figure 2.**
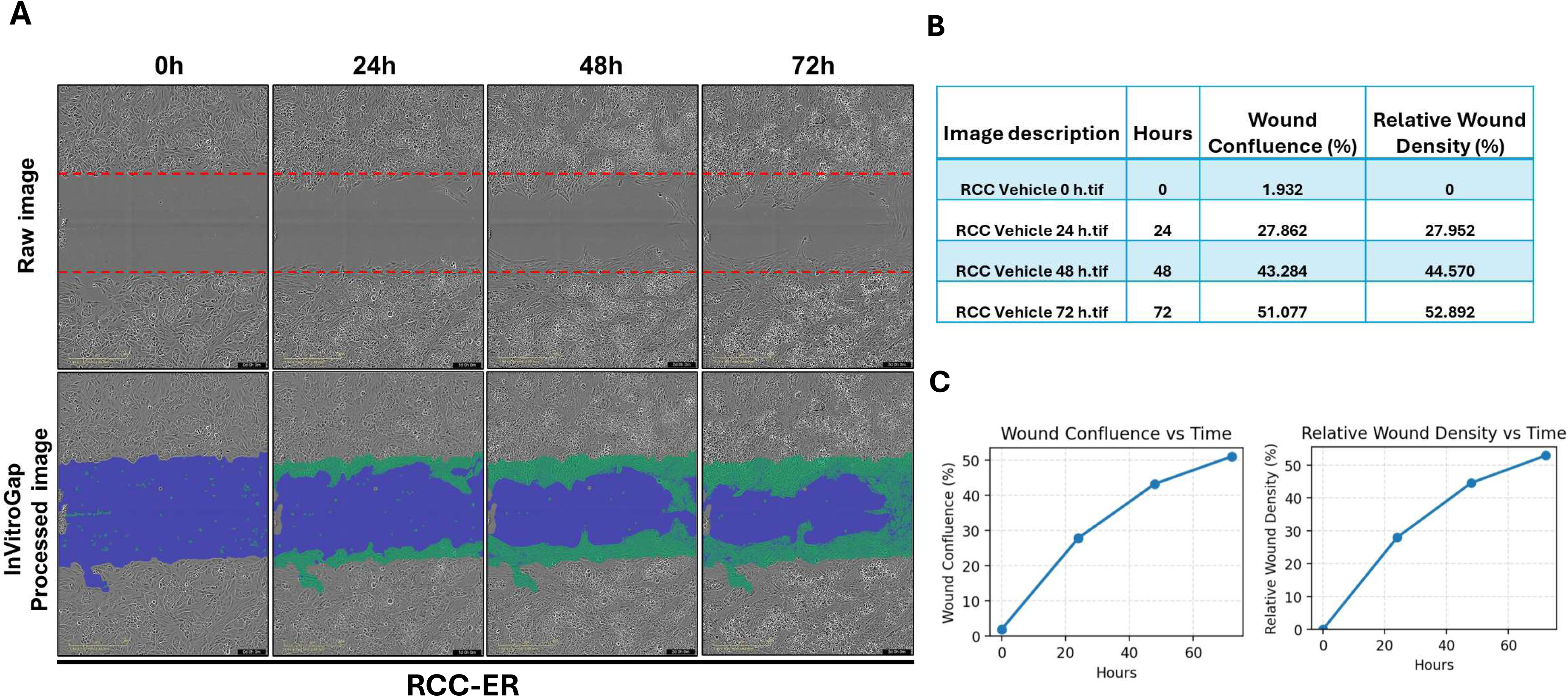
Representative InVitroGap analysis of wound-closure kinetics in RCC-ER cells. (**A**) Phase-contrast time-lapse images at 0, 24, 48, and 72 h (top row) with corresponding InVitroGap segmentation overlays (bottom row; blue = open wound area; green = cell-covered region). Scale bar = 400 µm. (**B**) Quantitative output table of wound confluence (%) and relative wound density (RWD, %) at each time point. (**C**) Time-course plots of wound confluence and RWD (mean ± SEM, *n* = 6 independent wells).

**Figure 3.**
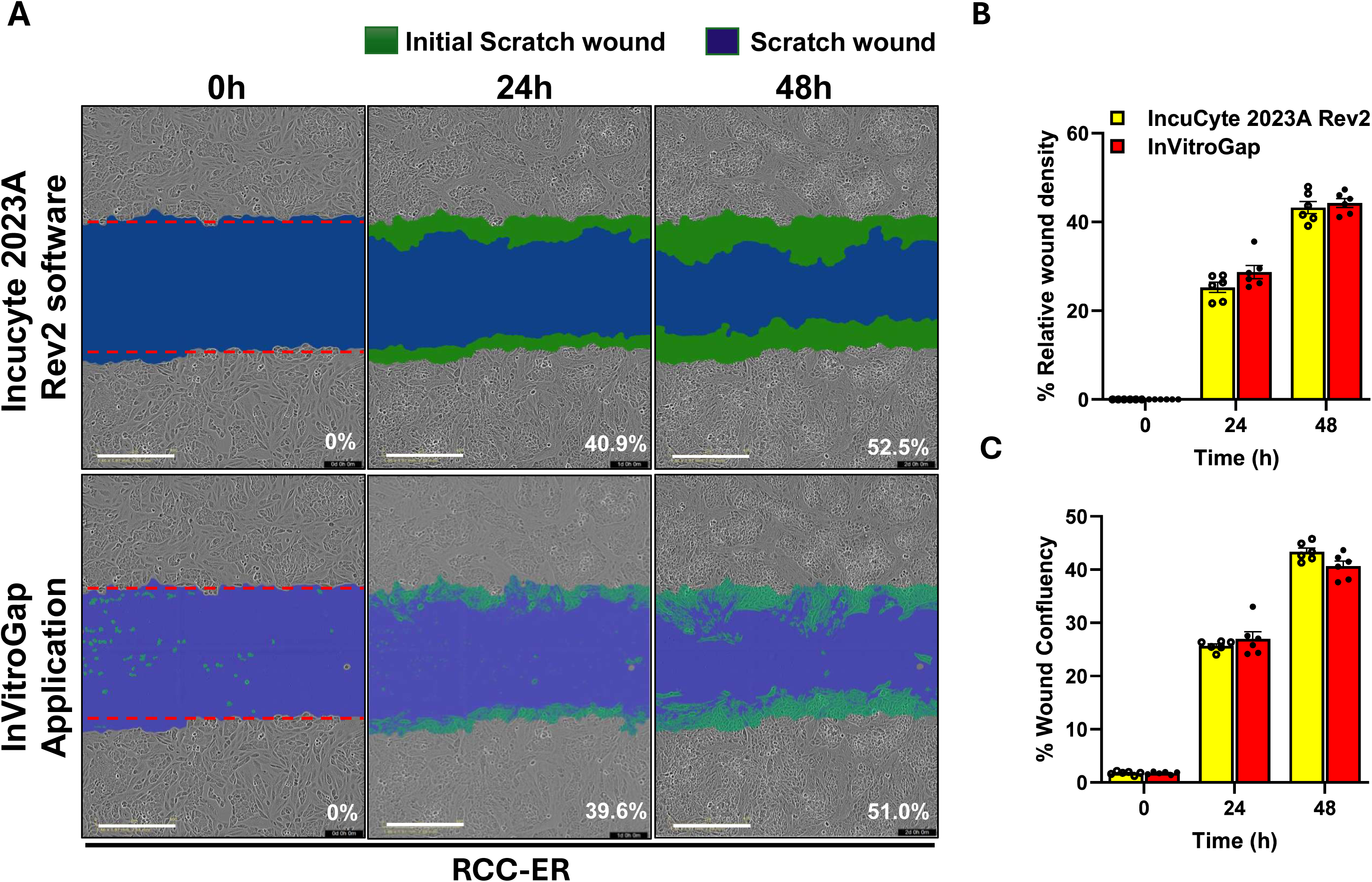
Benchmarking of InVitroGap against IncuCyte 2023A Rev2 in human RCC-ER cells. **(A)** Representative segmentation masks at 0, 24, and 48 h generated by IncuCyte 2023A Rev2 (top row) and InVitroGap (bottom row), with labeled wound confluence values. Scale bar = 400 µm. **(B)** Bar graphs comparing relative wound density (%) at the displayed time points. **(C)** Bar graphs comparing wound confluence (%) at the displayed time points. Data are presented as mean ± SEM (n = 6 wells; paired t-test, p > 0.05 for all displayed comparisons).

**Figure 4.**
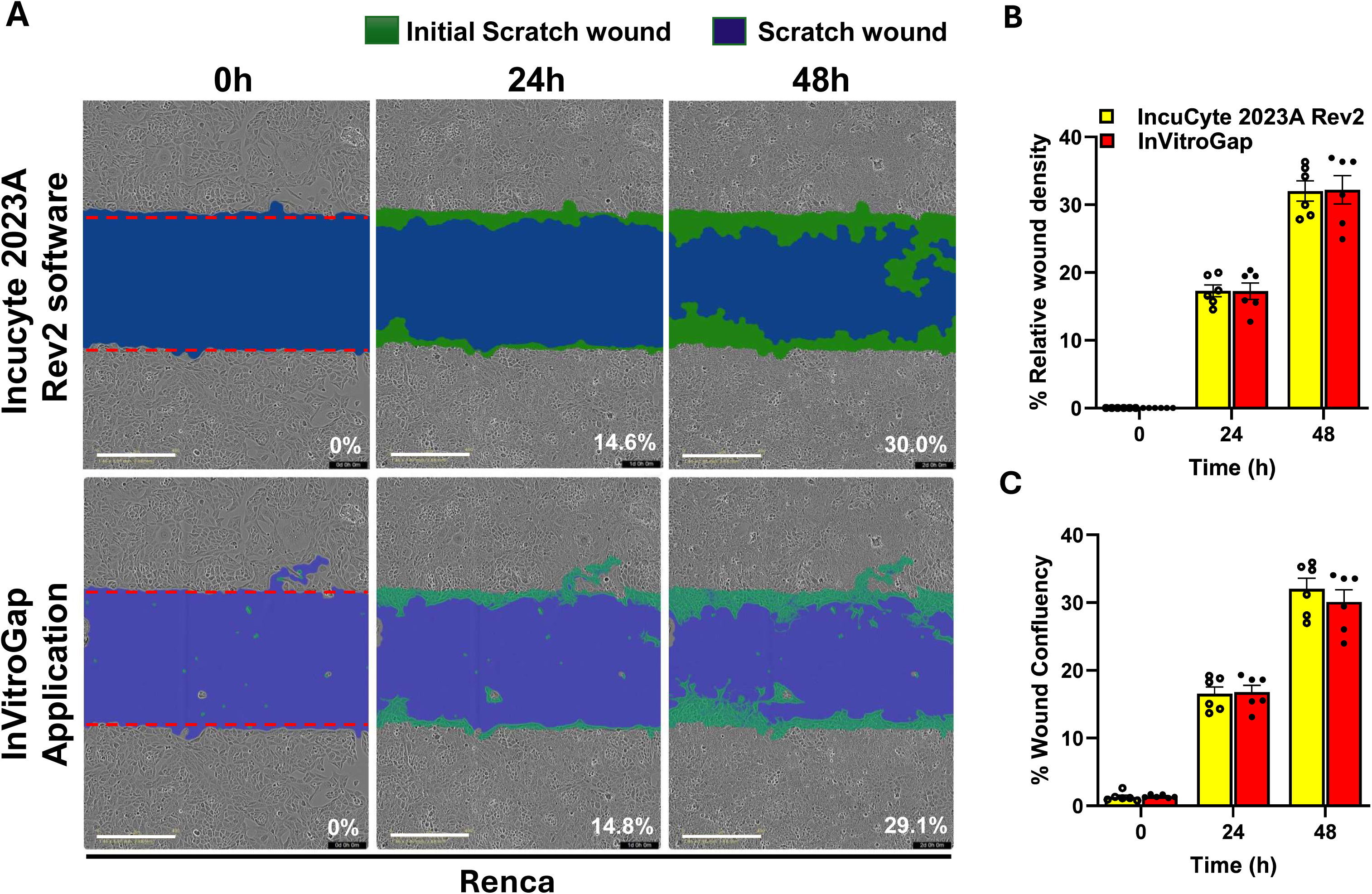
Benchmarking of InVitroGap against IncuCyte 2023A Rev2 in murine Renca cells. **(A)** Representative segmentation masks at 0, 24, and 48 h generated by IncuCyte 2023A Rev2 (top row) and InVitroGap (bottom row), with labeled wound confluence values. Scale bar = 400 µm. **(B)** Bar graphs comparing relative wound density (%) at the displayed time points. **(C)** Bar graphs comparing wound confluence (%) at the displayed time points. Data are presented as mean ± SEM (n = 6 wells; paired t-test, p > 0.05 for all displayed comparisons). High agreement was maintained despite distinct cell morphology compared with RCC-ER cells.

**Figure 5.**
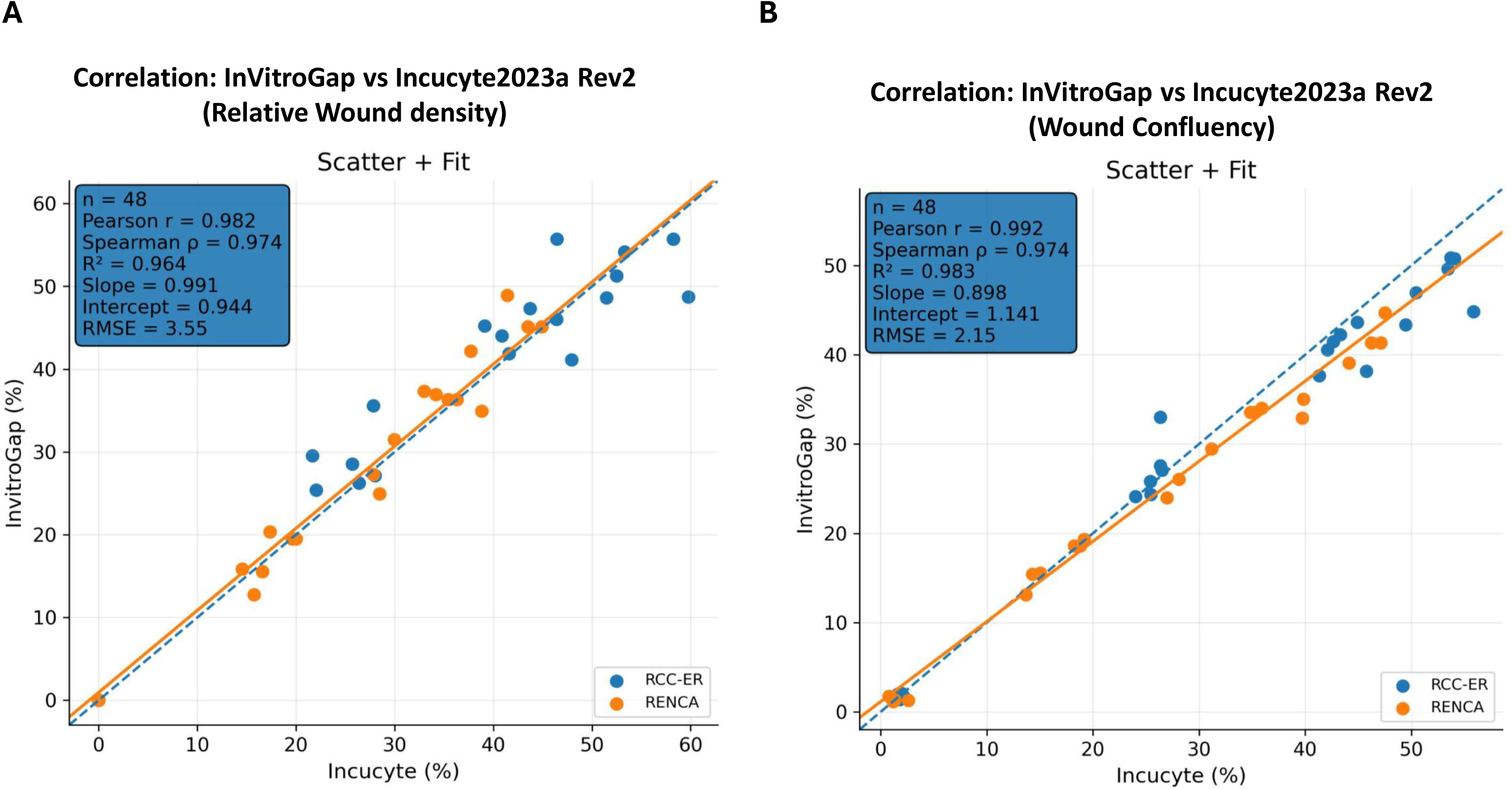
Pooled correlation scatter plots comparing InVitroGap and IncuCyte 2023A Rev2 outputs across all time points (0–72 h) for both cell lines combined. **(A)** Relative wound density (RWD): Pearson r = 0.982, Spearman ρ = 0.974, and R² = 0.964 (n = 48). **(B)** Wound confluence: Pearson r = 0.992, Spearman ρ = 0.974, and R² = 0.983 (n = 48). The identity line (y = x, dashed) is shown for reference. Data points cluster tightly along the identity line across the observed measurement range, confirming strong linear agreement between platforms.

**Figure 6.**
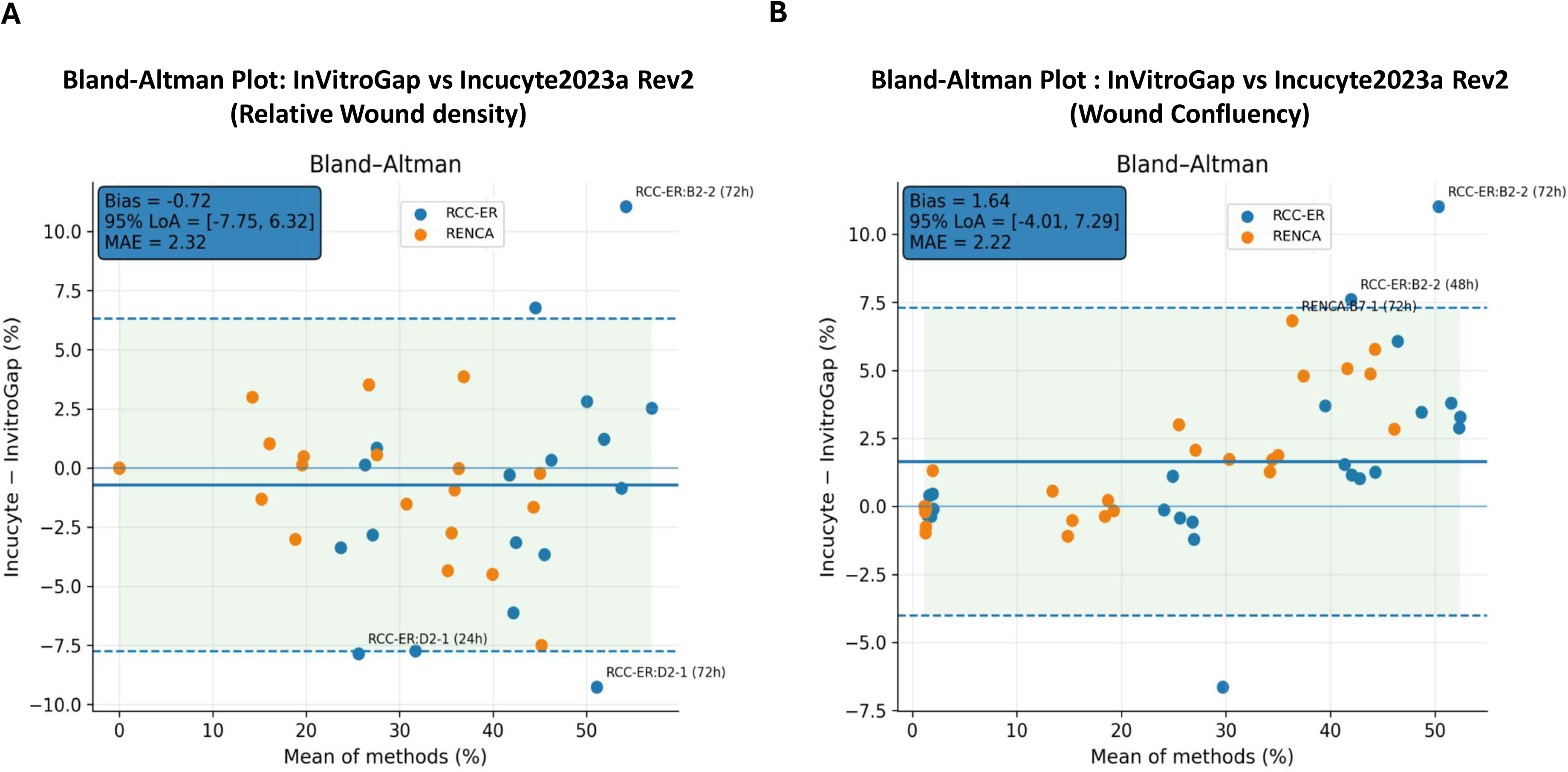
Pooled Bland–Altman agreement analysis comparing InVitroGap and IncuCyte 2023A Rev2. Differences are calculated as IncuCyte − InVitroGap and plotted against the mean of both measurements. **(A)** Relative wound density: mean bias = −0.72%, 95% limits of agreement = −7.75% to +6.32%, and MAE = 2.32% (n = 48). **(B)** Wound confluence: mean bias = +1.64%, 95% limits of agreement = −4.01% to +7.29%, and MAE = 2.22% (n = 48). Solid central lines indicate the mean difference; outer dashed lines indicate the 95% limits of agreement (bias ± 1.96 × SD). No strong proportional bias is apparent across the observed measurement range.

### 2.3. IncuCyte software analysis

Wound confluence was quantified using IncuCyte 2023A Rev2 software (Sartorius, Göttingen, Germany). Phase-contrast images were processed using a customized Analysis Definition to ensure accurate segmentation of the cell monolayer from the background. A segmentation adjustment of 1.6 was applied, with the threshold biased toward “Cells” to optimize edge detection. Post-segmentation cleanup included an Adjust Size of −2 pixels to refine mask boundaries, while the Hole Fill parameter remained at 0.0 µm². A minimum area filter of 500 µm² was implemented to exclude small debris and non-specific artifacts. No eccentricity or maximum area filters were applied, permitting the inclusion of all large-scale confluent regions within the wound area.

### 2.4. InVitroGap application and algorithm

InVitroGap is a freely available, open-source Python-based application with a browser-accessible interface at https://invitrogap.vercel.app/. It implements a computationally efficient, texture-based image-processing pipeline designed to identify wound regions and quantify wound-closure dynamics from standard phase-contrast time-lapse images without requiring proprietary software or hardware.

#### 2.4.1. Image pre-processing

All phase-contrast images were converted to 8-bit grayscale to ensure consistent feature extraction across sequential time points. Gaussian smoothing (σ = 1.0) was applied to suppress sensor noise while preserving high-frequency features associated with cell boundaries. Image texture was subsequently quantified using the Sobel gradient magnitude[17], exploiting the observation that confluent cell monolayers exhibit significantly higher textural variance than open wound regions.

#### 2.4.2. Wound region initialization and reference band definition

The wound region of interest (ROI) was defined exclusively from the baseline image (*t*₀) and maintained as a fixed spatial reference throughout the entire time series. Pixels with gradient magnitudes below the *p*th percentile (*p* = 30–40) were provisionally classified as wound area. This binary mask was refined using morphological closing and opening operations (disk radius = 2–3 pixels), followed by removal of small disconnected components. The largest remaining connected region was designated as the wound ROI.

To establish a local texture baseline for adaptive segmentation, a reference band was generated by dilating the wound ROI by 50 pixels and subtracting the original wound mask. This annular reference band captures the fully confluent monolayer immediately adjacent to the scratch and provides an experiment-specific texture baseline for cell detection across all subsequent time points.

#### 2.4.3. Adaptive segmentation and time-series analysis

For all subsequent frames (*t* = 24, 48, and 72 h), segmentation thresholds were dynamically defined as the *q*th percentile of gradient magnitudes within the reference band (default *q* = 5), ensuring robustness to illumination and contrast variability across experiments. Pixels exceeding this adaptive threshold within the fixed wound ROI were classified as cell-covered, representing migrating cells contributing to wound closure. Connected components smaller than 80 pixels were excluded to suppress floating debris and imaging artifacts. The complete four-timepoint analysis was performed in approximately 10 seconds on a standard desktop computer.

#### 2.4.4. Quantitative metrics

Wound-healing progression was quantified using two established confluence-based metrics. Wound confluence was defined as the percentage of the wound ROI occupied by cell-covered pixels at each time point. Relative wound density (RWD) was calculated to normalize cell detection against changes in density in the reference region using the equation:

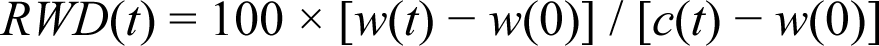

where *w*(*t*) and *w*(0) denote wound region cell densities at time *t* and baseline, respectively, and *c*(*t*) represents the mean cell density within the reference band at time *t*.

### 2.5. Statistical analysis

All data are presented as mean ± standard error of the mean (SEM) from n = 6 wells per cell line from a single representative experiment. Because the same wells were measured repeatedly over time, pooled image-level analyses were treated as benchmarking analyses rather than formal equivalence testing. Agreement between IncuCyte and InVitroGap was assessed using pooled image-level data from RCC-ER and Renca cells across all time points (0, 24, 48, and 72 h; total n = 48 observations per metric). For wound confluence and relative wound density (RWD), scatter plots were generated with IncuCyte values on the x-axis and InVitroGap values on the y-axis, together with the identity line (y = x) and an ordinary least squares regression fit. Pearson’s correlation coefficient (r), Spearman’s rank correlation coefficient (ρ), and the coefficient of determination (R²) were calculated. Method agreement was further evaluated by Bland–Altman analysis, using the difference between methods (IncuCyte − InVitroGap), with calculation of mean bias[18, 19], 95% limits of agreement, and mean absolute error (MAE). Statistical analyses were performed in Python using pandas, scipy.stats, and matplotlib. No formal equivalence margin was prespecified. Raw data and analysis scripts are available from the corresponding author upon reasonable request.

## Declaration of generative AI and AI-assisted technologies in the manuscript preparation process

During the preparation of this work the author RKA used ChatGPT in order to paraphrase some part of the introduction and discussion. After using this tool/service, the author reviewed and edited the content as needed and take(s) full responsibility for the content of the published article.

## 3. Results

### 3.1. InVitroGap workflow and image analysis pipeline

The InVitroGap application features a streamlined GUI designed to automate the quantification of wound-healing kinetics from time-lapse phase-contrast images (Figure 1B). The processing pipeline establishes a fixed wound ROI at *t* = 0 h and constructs an adaptive reference band that captures the confluent monolayer immediately adjacent to the scratch, providing an experiment-specific baseline for segmentation at all subsequent time points (Figure 1A). This architecture, combined with Sobel gradient-based texture extraction and dynamic percentile thresholding, enables high-fidelity measurement of wound confluence and RWD across the full-time course (Figures 2B and 2C).

In this study, analysis parameters were configured as follows: Gaussian blur σ = 1.00; wound edge smoothing = 2–3 pixels; wound smoothness percentile = 30–40; reference band thickness = 50 pixels; minimum cell object size = 80 pixels; minimum wound size = 500 pixels (Figure 1B). These settings were applied consistently across all image sets. The automated analysis successfully tracked cellular invasion into the cell-free area over the 72-hour observation period (Figure 2A). Segmentation masks accurately delineated wound boundaries, capturing progressive closure from an initial wound confluence of 1.93% at 0 h to 51.08% by 72 h (Figures 2B and 2C).

### 3.2. Validation against IncuCyte in human RCC-ER cells

To validate the accuracy of InVitroGap, outputs from identical time-lapse image sets were compared with those of IncuCyte 2023A Rev2 software using human RCC-ER cells across four time points (0, 24, 48, and 72 h). For visual clarity, the representative image panels and per-time-point summary plots shown in Figure 3 are limited to 0, 24, and 48 h; however, 72 h measurements from the same dataset were retained in the pooled agreement analyses presented in Figures 5 and 6. High concordance in segmentation mask generation was observed between platforms (Figure 3A). At 48 h, IncuCyte reported wound confluence of 52.5% compared with 51.0% for InVitroGap, a difference of 1.5 percentage points (Figure 3A). Quantitative analysis of both RWD (Figure 3B) and wound confluence (Figure 3C) confirmed no statistically significant differences between the two platforms at the displayed time points (p > 0.05), with both tools tracking nearly identical migration trajectories.

### 3.3. Validation against IncuCyte in murine Renca cells

To assess generalizability across cell types, validation was extended to Renca, a murine renal adenocarcinoma cell line with distinct cellular morphology relative to RCC-ER cells, including smaller cell size and more compact colony structure. For visual clarity, the representative image panels and per-time-point summary plots shown in Figure 4 are limited to 0, 24, and 48 h; however, 72 h measurements from the same dataset were retained in the pooled agreement analyses presented in Figures 5 and 6. Despite these morphological differences, InVitroGap maintained high concordance with IncuCyte segmentation masks (Figure 4A). At 48 h, IncuCyte reported wound confluence of 30.0% compared with 29.1% for InVitroGap (Figure 4A). Statistical analysis confirmed that RWD (Figure 4B) and wound confluence (Figure 4C) were not significantly different between platforms at the displayed time points (p > 0.05), with overlapping data trajectories and minimal inter-tool variance.

### 3.4. Performance under non-standard experimental conditions

To further assess the versatility of InVitroGap, its performance was evaluated on crisscross scratches generated manually with 200 µL pipette tips and imaged using standard brightfield microscopy rather than the IncuCyte SX5 system (Supplementary Figure S1). This configuration simultaneously tested two important edge cases: an irregular, non-linear wound geometry and a lower-specification imaging environment without automated focus control.

Despite these challenges, InVitroGap successfully identified and delineated intersecting wound boundaries (Supplementary Figure S1A). The algorithm tracked progressive closure from 0% confluence at baseline to 37.47% at 48 h (Supplementary Figure S1C), demonstrating that the adaptive reference-band strategy remains effective even when wound geometry deviates substantially from the standard linear form. These results confirm that InVitroGap is applicable to data from standard laboratory microscopes and is not restricted to IncuCyte-acquired images.

### 3.5. Statistical correlation and agreement analysis

To provide a quantitative assessment of agreement between InVitroGap and IncuCyte 2023A Rev2, Pearson and Spearman correlation analyses and Bland–Altman plots were generated using pooled image-level data across both cell lines and all time points (0–72 h; total n = 48 observations per metric). As noted in the Methods, these pooled analyses should be interpreted as benchmarking summaries because repeated measurements from the same wells were included over time.

Scatter plots demonstrated strong pooled linear correlations (Figure 5). For RWD, the pooled comparison across both cell lines yielded Pearson r = 0.982, Spearman ρ = 0.974, and R² = 0.964. For wound confluence, the pooled comparison yielded Pearson r = 0.992, Spearman ρ = 0.974, and R² = 0.983. In both analyses, data points clustered tightly along the line of identity (y = x, dashed), supporting close agreement between InVitroGap and IncuCyte across the observed measurement range.

Bland–Altman analysis quantified systematic bias and limits of agreement (Figure 6). Differences were calculated as IncuCyte minus InVitroGap. For RWD (Figure 6A), the mean bias was −0.72%, with 95% limits of agreement from −7.75% to +6.32% and MAE = 2.32%. For wound confluence (Figure 6B), the mean bias was +1.64%, with 95% limits of agreement from −4.01% to +7.29% and MAE = 2.22%. Together, these findings indicate low average inter-method discrepancy and close agreement between platforms across the observed measurement range within the tested datasets. Visual inspection of the Bland–Altman plots did not suggest meaningful proportional bias, and the great majority of observations fell within the 95% limits of agreement.

## 4. Discussion and Conclusion

We present InVitroGap, an open-source tool for automated, time-resolved quantification of wound closure in in vitro scratch assays. When benchmarked against IncuCyte, InVitroGap produced wound confluence and relative wound density (RWD) measurements that closely matched the commercial reference within the tested datasets (pooled R² = 0.964 for RWD and R² = 0.983 for wound confluence; absolute bias ≤ 1.64%). These results support the utility of the tool for comparative and longitudinal studies of wound closure.

A key feature of InVitroGap is its adaptive segmentation, which adjusts to each experiment rather than using fixed thresholds. The algorithm first identifies the wound at the starting time point and defines boundaries using a local reference region of the surrounding confluent monolayer. This approach allows the method to handle variations in illumination, contrast, and cell density over time, and accurately track wound edges even during later stages of closure when cells begin to grow back. By focusing on local, biologically relevant information, InVitroGap achieves consistent and precise segmentation across diverse experimental conditions and image qualities.

We observed a small positive mean bias for wound confluence when differences were expressed as IncuCyte minus InVitroGap, indicating that IncuCyte tended to report slightly higher confluence values than InVitroGap on average. This likely reflects the fact that InVitroGap employs a gradient-based texture metric with adaptive percentile thresholding, whereas the commercial reference uses a proprietary, hardware-optimized pipeline. These methodological differences may lead to minor systematic discrepancies; however, the relative trends over time and between experimental conditions were preserved, supporting reliable comparative and longitudinal analyses.

The framework also demonstrated robustness to biological and technical variability. Agreement with the reference system was maintained across both RCC-ER and Renca cell lines, despite differences in morphology, growth patterns, and wound closure kinetics. In addition, InVitroGap successfully quantified closure in irregular, manually generated crisscross wounds imaged with standard brightfield microscopy, highlighting its versatility. Its independence from specific imaging platforms or standardized wound configurations makes the tool particularly suitable for laboratories with heterogeneous equipment or non-standard assay setups.

From a practical standpoint, InVitroGap combines computational efficiency with user control. Time-series images can be processed rapidly on standard desktop hardware, allowing the analysis of multiple conditions and replicates without bottlenecks. Automated segmentation reduces operator-dependent variability, while a graphical interface enables users to inspect and fine-tune parameters when necessary. This combination of automation and user oversight ensures both high throughput and quality control.

Another advantage of the framework is its transparency and reproducibility. InVitroGap uses deterministic image-processing steps and user-adjustable parameters, allowing the same image series to be re-analyzed consistently across experiments. Raw and processed outputs can be reviewed together with segmentation overlays, facilitating quality control and iterative parameter refinement in quantitative scratch-assay studies.

Some limitations remain. Current benchmarking focused on two cell lines, a modest number of wells, and one representative experiment, and additional studies will be required to confirm performance across broader biological systems and imaging conditions. In addition, the pooled agreement analyses included repeated measurements from the same wells over time and therefore should be interpreted as benchmarking summaries rather than formal equivalence testing. InVitroGap is currently optimized for two-dimensional phase-contrast and brightfield imaging; its performance in fluorescence or three-dimensional imaging modalities has yet to be determined. Future development should include validation across independent experiments, additional imaging systems, and broader assay conditions.

In conclusion, InVitroGap provides a robust and reproducible framework for automated scratch-assay analysis. By integrating adaptive, context-sensitive segmentation with efficient processing and methodological transparency, it addresses key limitations of existing approaches while remaining independent of proprietary hardware. Its versatility and accessibility make it suitable for a wide range of experimental conditions, and continued development should expand its applicability to more complex imaging scenarios and higher-throughput workflows, supporting standardized quantification of wound closure in biomedical research.

## CRediT Author Contribution Statement

R.K.A. contributed to conceptualization, methodology, investigation, formal analysis, and writing of the original draft, as well as review and editing of the manuscript. M.S. contributed to software development, statistical formal analysis, and manuscript review and editing. S.R.D. performed investigation related to the crisscross assay experiments. C.J.W. provided supervision. L.B. contributed to conceptualization, project administration, supervision, and manuscript review and editing.

## Data Availability Statement

The raw image data and analysis scripts supporting the findings of this study are available from the author (L. Bukavina) or (R. Arya) upon reasonable request.

## Software Availability Statement

InVitroGap is freely available as a Python-based application with a browser-accessible interface at https://invitrogap.vercel.app/. The application requires no installation and is compatible with all major operating systems and modern web browsers. There are no hardware, licensing, or subscription requirements.

## Acknowledgements

The authors thank Dr. Philip Abbosh MD, PhD (Fox Chase Cancer Center, PA, USA) for generously providing the RCC-ER and Renca cell lines used in this study. The authors also thank the Cleveland Clinic institutional core facilities for supporting cell culture and imaging infrastructure.

---

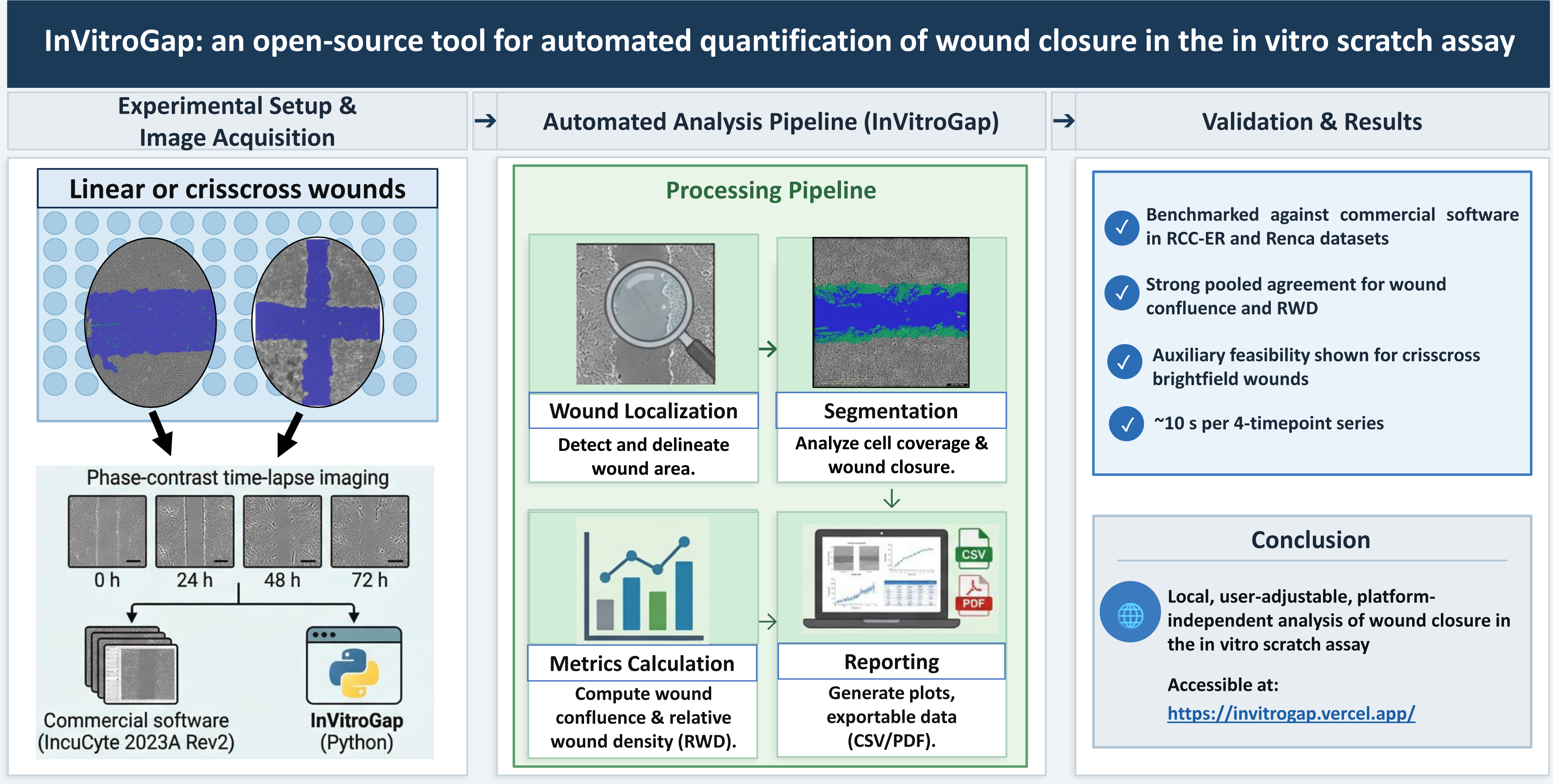

**Supplementary Figure S1.**
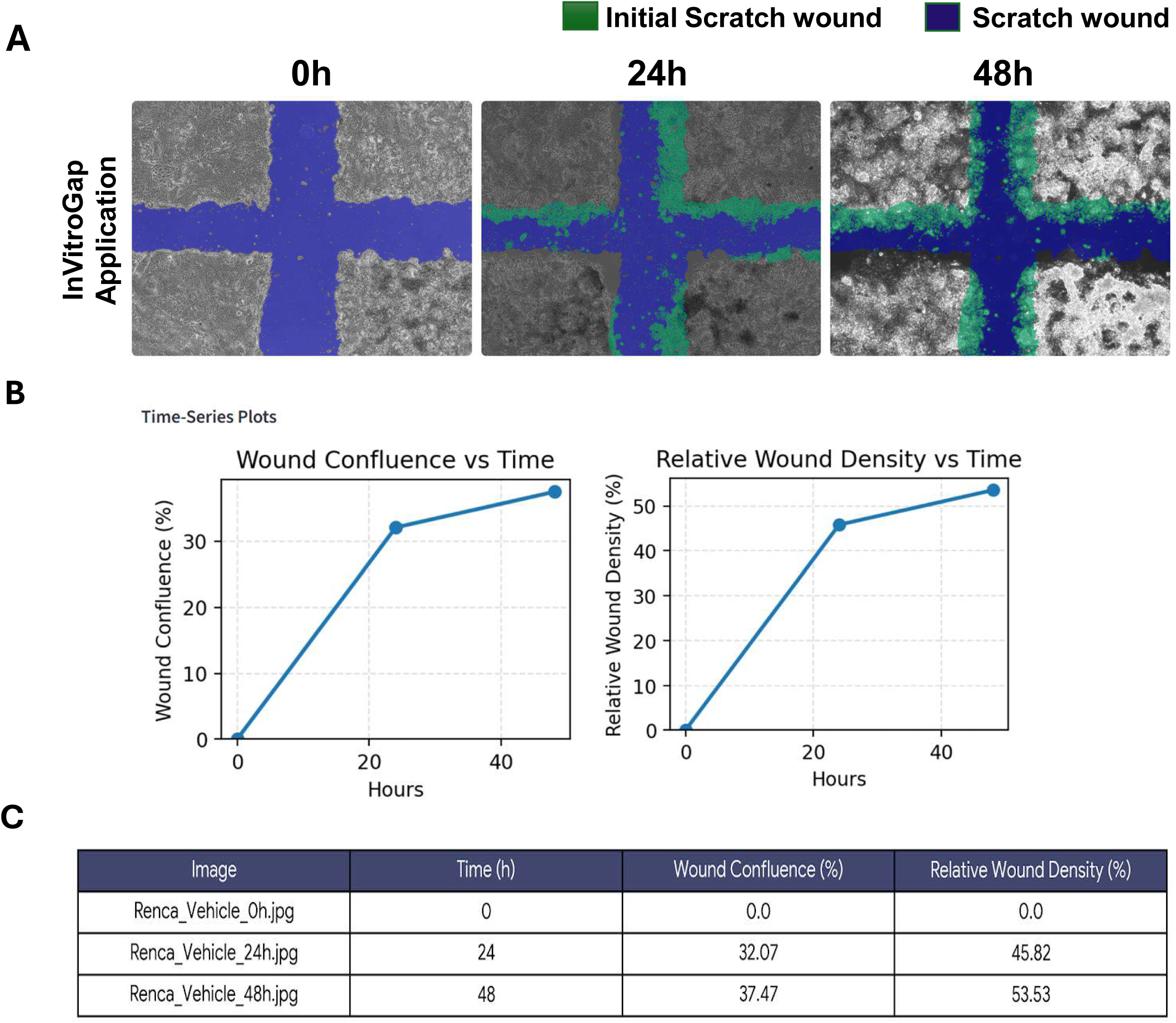
InVitroGap performance on non-standard experimental conditions. (**A**) Brightfield time-lapse images of manual crisscross wounds (200 µL pipette tip) at 0, 24, and 48 h with corresponding InVitroGap segmentation overlays (blue = open wound area; green = cell-covered region). The algorithm successfully identifies both intersecting wound boundaries within a single ROI. Scale bar = 400 µm. (**B**) Time-course plots of wound confluence and RWD for the crisscross wound configuration. (**C**) Quantitative output table confirming reliable wound tracking (0% to 37.47% confluence at 48 h) despite irregular wound geometry and brightfield imaging conditions.

**Supplementary Table 1.**
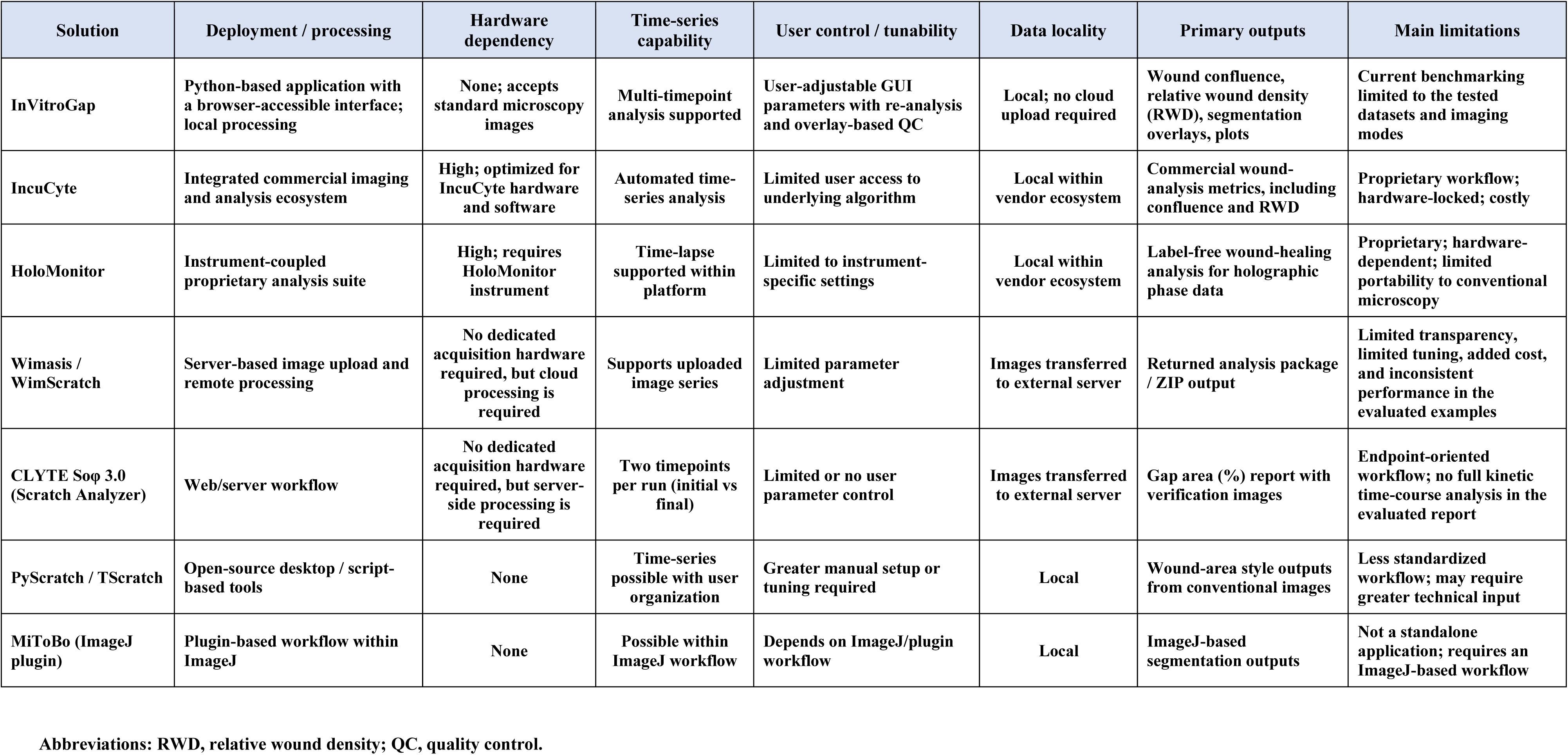
Comparison of InVitroGap with representative scratch-assay image-analysis solutions. Comparison of InVitroGap with representative commercial, cloud-based, and open-source scratch-assay image-analysis platforms, including IncuCyte, HoloMonitor, Wimasis/WimScratch, CLYTE Soφ 3.0, PyScratch/TScratch, and MiToBo. Solutions are compared with respect to deployment/processing workflow, hardware dependency, time-series capability, user control/tunability, data locality, primary outputs, and major limitations. This table is intended as a workflow- and feature-level comparison rather than a direct head-to-head performance benchmark on a shared dataset. Abbreviations: RWD, relative wound density; QC, quality control.

